# Modelling the interplay between inadvertent social information use and a pesticide-induced foraging bias in bumblebees’ crop visitation

**DOI:** 10.1101/2025.07.29.667368

**Authors:** Zoltán Tóth

## Abstract

Understanding the drivers of patch visitation in agricultural landscapes is essential for maintaining pollinator-mediated agroecosystem services, yet the extent to which behavioural mechanisms influence pollinators’ visitation rates has received little attention. Using an individual-based modelling framework (BEE-STEWARD), I examined how inadvertent social information (ISI) use and a recently documented pesticide-induced foraging bias influence nectar and pollen visitation rates of buff-tailed bumblebees (*Bombus terrestris*) to crop patches located near or far from the colonies. Simulations showed that ISI use primarily increased nectar visitation to all crop patches and pollen visitation to nearby crop patches, whereas foraging bias increased pollen visitation to all crop patches and nectar visitation to nearby crop patches. However, the two behavioural parameters had no interactive effects on crop visitation rates during nectar or pollen foraging. Despite the pronounced shifts in visitation patterns, total resource collection and colony demographics, including colony growth and reproductive output, remained largely unaffected. Overall, results indicate that ISI use and foraging biases are important drivers of spatial foraging patterns in bumblebees. Incorporating such behavioural pathways into pollinator foraging models may improve predictions of spatial visitation patterns and resource exploitation across agricultural landscapes, with potential implications for pesticide risk assessments.

## Introduction

Social information provided by conspecifics or heterospecifics (Danchin et al. 2004; Seppänen et al. 2007) has a crucial role in adaptation to predation risk (Coolen et al. 2005; Martín et al. 2006; Pays et al. 2013; Cruz et al. 2020), human disturbance (Banks et al. 2013; Schmidt et al. 2015; Gruber et al. 2019), or resource exploitation (Laidre 2010; Webster and Laland 2017; Pouca et al. 2020; Curk et al. 2025) in a wide variety of animals. Social information can originate either from evolved, actively produced signals, or from inadvertently produced cues. Inadvertently produced cues (visual, acoustic, chemical, etc.) include the presence, routine behaviour, or the product of the behaviour of others that convey information to the observers about the environment or the physiological state of the cue-producing individual (Gil et al. 2018; Tóth et al. 2020). Unlike social learning, inadvertent social information (ISI) use is not characterised by the presence of a conditioning phase (Leadbeater 2015), and does not necessarily result in the acquisition of new knowledge or skill. ISI use, nevertheless, often generates strong correlations in the behaviours and space use of multiple individuals (Laundré et al. 2010; Gil et al. 2017), making the social environment capable of shaping population dynamics and even ecosystem structure and function (Parejo and Avilés 2016; Gil and Hein 2017; Gil et al. 2018, 2019). ISI use can also contribute to the spread of maladaptive behaviours, potentially accelerating population declines and increasing extinction risk through information-mediated Allee effects (Kokko and Sutherland 2001; Donaldson et al. 2012; Sigaud et al. 2017; Wilson et al. 2020).

ISI use is a well-known phenomenon in pollinating insects (Grüter and Leadbeater 2014). Bumblebees, like many other bee species (Yokoi et al. 2007; Wray et al. 2012), often rely on social cues when they do not have personal information about surrounding food sources or finding them would be more costly (Kawaguchi et al. 2006; Jones et al. 2015).

These pollinators possess remarkable cognitive abilities, and social learning has been demonstrated in a variety of contexts (e.g., Loukola et al. 2017). However, the possibility of social learning does not preclude ISI use and several studies have shown its fundamental role in bumblebees’ foraging (e.g., Baude et al. 2008; Leadbeater and Florent 2014; Smolla et al. 2016). Under natural conditions, ISI use is often advantageous as it facilitates the localisation and exploitation of new food sources (Worden and Papaj 2005; Avarguès-Weber and Chittka 2014) and helps to avoid predators (Dawson and Chittka 2014). The presence of conspecifics can also provide a cue about the quantity of resources in the visited flowers (Leadbeater and Chittka 2009). However, relying on social information may also limit the time spent searching for alternative food sources and making independent foraging decisions, resulting in suboptimal foraging (Avarguès-Weber et al. 2018). In agricultural habitats, the attraction of conspecifics may contribute to the higher exploitation of crop patches, which could have consequences for several ecological processes, including exposure to pesticide contamination. Such exposure has been associated with impaired foraging behaviour or physiology and, ultimately, reduced colony growth or fitness (e.g., Baron et al. 2017; Crall et al. 2019). Interestingly, the effects of ISI use on crop visitation patterns are largely understudied in most pollinator species and remain unaddressed in recent theoretical models (e.g., Blasi et al. 2022; Carturan et al. 2023; Lonsdorf et al. 2024).

Spatial variation in resource quality and availability can strongly influence pollinator foraging decisions, yet the behavioural processes shaping patch visitation patterns are not fully understood. For instance, evidence suggests that exposure to pesticides can alter individual foraging behaviour. Such behavioural biases may manifest as a preference for pesticide-treated food sources (Kessler et al. 2015; Arce et al. 2018), impaired cognitive abilities and feeding motivation (Siviter and Muth 2022; Gray et al. 2024), decreased resource collection (e.g., Lämsä et al. 2018; Muth and Leonard 2019), or a reduced ability to orient toward a reward-associated floral scent source (Kárpáti et al. 2024). These behavioural changes may, in turn, alter visitation patterns across landscapes, potentially affecting the relative use of different habitat types, including those that receive – directly or indirectly (Ward et al. 2022) – regular pesticide treatments (Pohorecka et al. 2012; Byrne et al. 2014). However, the potential contribution of such behavioural pathways to spatial foraging patterns in pollinators is largely unexplored.

In this study, I extended the existing individual-based model of BEE-STEWARD (Twiston-Davies et al. 2021) to explore how ISI use and a recently documented pesticide-induced foraging bias influence crop patch visitation rates in buff-tailed bumblebees (*Bombus terrestris*) in a simulated agricultural landscape. This species is an important pollinator of both crops and wildflowers in temperate regions (Hutchinson et al. 2021), and a common visitor of the mass-flowering oilseed rape (*Brassica napus*; ‘OSR’ henceforward), a dominant oil crop in Europe (Perrot et al. 2024). By including the possibility of ISI use among foragers and reduced ability to locate rewarding food sources as a foraging bias (following Kárpáti et al. 2024) in the model, I predicted that bumblebees would exploit the available food sources more efficiently in the presence of ISI use, leading to a shift in patch visitation patterns and the collection of more resources compared to the original model setting. In the presence of foraging bias, I expected visitation patterns to shift and foraging efficiency to decrease. This effect could also be amplified by ISI use via the attraction of other foragers to suboptimal resource patches. In this study, I simulated the attraction of conspecifics to the same resource patches, which does not involve the acquisition of new knowledge (i.e., it is not social learning) or the exploitation of evolved signals (i.e., it is not communication), but falls within the definition of ISI use (i.e., the exploitation of social cues originating from the routine behaviour of others).

## Materials and methods

### Description of the applied model

Simulations were run in BEE-STEWARD (www.beesteward.co.uk; Twiston-Davies et al. 2021). This simulation tool incorporates landscape data and the full life cycle of bumblebees, allowing researchers to test the impact of land-use characteristics and environmental stressors on colony and population dynamics in various bumblebee species. Individual behaviour is determined by stimuli and thresholds that scale up to colony-level processes, enabling the simulation of population survival and performance at the landscape level over multiple years. The underlying modules of the software are parameterised with empirical data from the UK, describing colony dynamics of six bumblebee species and plant species characteristics, including details on flower shape, resource production, and flowering period. The model also incorporates energy budgets and depletion of resources to facilitate the emergence of competition both within and between pollinator species. With sensitivity analysis and validation, the incorporated Bumble-BEEHAVE model has been demonstrated to make realistic predictions for buff-tailed bumblebee colonies within the boundaries of independent empirical observations (Becher et al. 2018). For a detailed description of the underlying models and procedures, see the supplementary materials of Becher et al. (2016, 2018) (Appendices S02 and S05, respectively).

In short, the landscape is 2-dimensional and comprises patches that provide foraging and/or nesting opportunities. Patches are spatially explicit and implemented as one or more food sources, each representing a single plant species. If a patch contains multiple food sources, these can be layered, i.e., placed at the same location and linked with each other. The simulations start at the beginning of the year with an initial number of hibernating queens that, after emerging from hibernation, search for suitable nest sites in semi-natural habitats (exact dates drawn from a characteristic distribution). Queens that survived the nest-searching process and successfully founded a colony must collect resources to lay the first batch of eggs, then forage and care for the brood simultaneously until the first adult workers emerge. After that, the queen’s activity is limited to egg laying, while the workers are responsible for nursing and foraging. Foragers’ choices on which patch to exploit are based on maximising the foraging rate for pollen and the energetic efficiency for nectar, primarily determined by the distance from the colony, handling time, and the amount of resource available in the patch (Knapp et al. 2019). Colony growth depends on resource collection, whereas population dynamics result from the number of mated queens that leave the colony and enter into hibernation until the following spring (Becher et al. 2018).

### Simulated landscape with specific foraging patches

The simulated landscape, consisting of 300 × 210 grid cells, contains one grassland and eight oilseed rape (*Brassica napus*) patches (Fig. 1). The central grassland patch provide nectar and pollen for the bumblebees’ year-around and also serves as a nesting habitat where queens can establish new colonies (randomly distributed within the grassland patch) after emerging from winter hibernation (Fig. S1). This patch is a good-quality habitat for buff-tailed bumblebees and contains 22 different flowering plant species (Becher et al. 2018). The OSR patches represent habitats that are unsuitable for nesting and solely consist of oilseed rape crop that flowers from day 114 until day 136 in each simulation year. All patches have a 200 m radius, resulting in a 125 663.7 m^2^ patch area. Due to the spatial distribution of OSR patches around the grassland patch, four OSR patches are closer to the central patch (distance between patch edges: 15 m) than the diagonally located OSR patches (186.9 m). The former patches are termed ‘close OSR patches’, whereas the latter ‘far OSR patches’; the exploitation of close and far OSR patches was examined separately.

**Figure 1.**
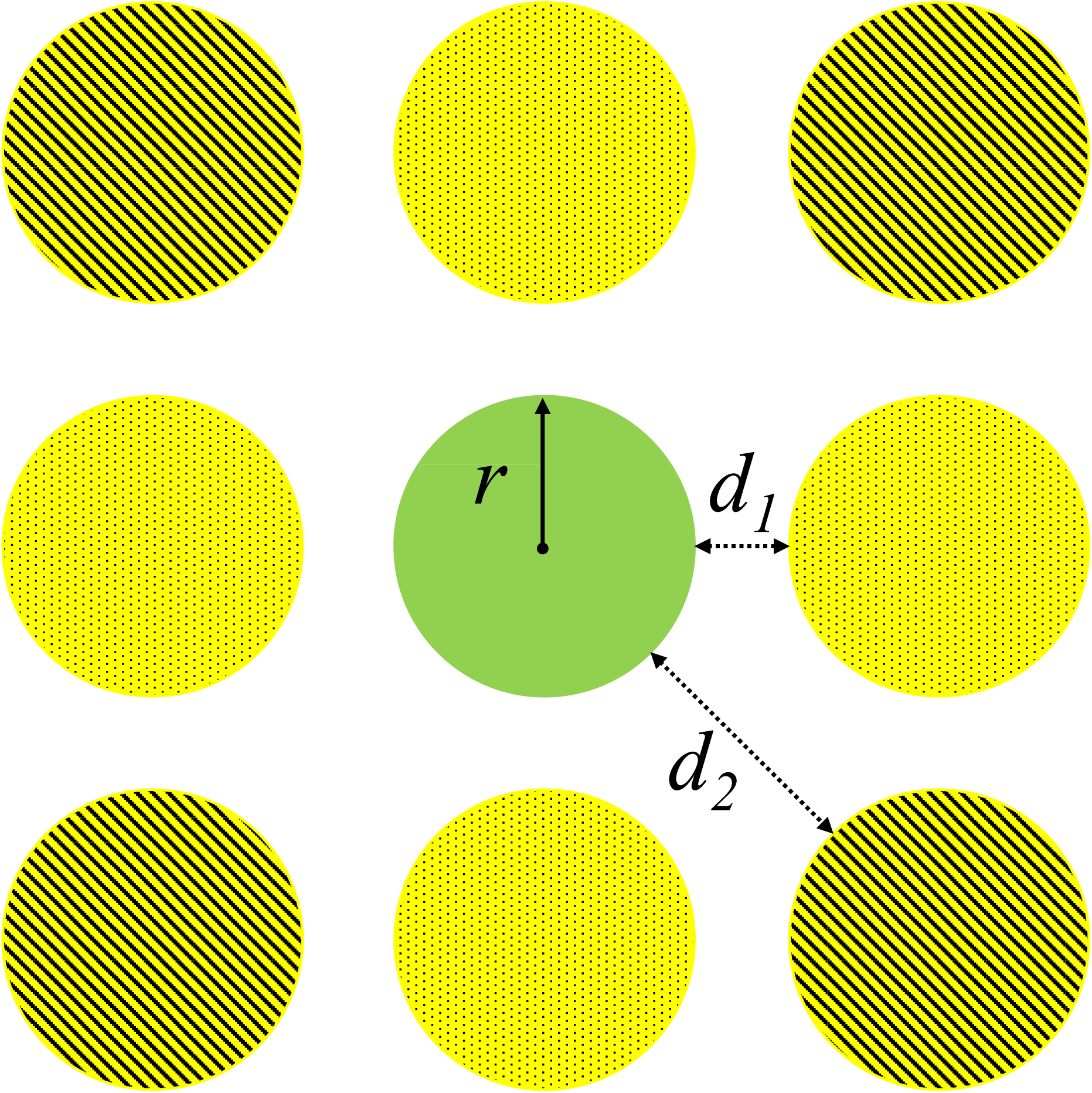
Schematic diagram of the simulated landscape. The simulated landscape consisted of a grassland (green) and multiple oilseed rape (OSR) patches (yellow). All patches had a 200 m radius (*r*), resulting in a 1256.64 m^2^ patch area. Due to the spatial distribution of OSR patches around the grassland patch, four OSR patches were closer to the central patch (*d*_1_: 15 m) than the diagonally located OSR patches (*d*_2_: 186.9 m). The former patches were termed ‘close OSR patches’ (with dotted pattern), whereas the latter ‘far OSR patches’ (with diagonal pattern). Distances on the diagram are not on scale. Alt text for Figure 1. Schematic of a simulated landscape with a central grassland patch surrounded by eight equally sized oilseed rape patches (all with 200 m radius). Four ‘close’ patches are positioned 15 m from the grassland, while additional ‘far’ patches are located diagonally at 186.9 m. Close and far patches are visually distinguished by different patterns.

### Model extensions

I modified the original code by incorporating the parameters ‘ISI use’ and ‘foraging bias’ into the model (Table 1). The model does not explicitly simulate pesticide exposure level as it would require detailed information on pesticide concentrations in nectar and pollen and their temporal dynamics in treated crops, which are still poorly characterized for many pesticides. Instead, it simulates two behavioural pathways that may influence patch visitation patterns, which, in turn, have been shown to be positively related to pesticide exposure risk in several empirical studies (e.g., Pohorecka et al. 2012; Byrne et al. 2014; Arce et al. 2018). I also included additional functions in the extended version to obtain patch-specific nectar and pollen visitation data. To reduce the computational time of the simulations, the code of the original model was converted from NetLogo 5.3.1 to 6.4.0 (Wilensky 1999).

**Table 1.** Additional parameters in the extended BEE-STEWARD model.

| **Parameter** | **Description** | **Range**<br>**(unit)** |
| --- | --- | --- |
| ISI use | When using ISI, foragers collect nectar or pollen from the closest patches where conspecifics are also present, provided such foraging sites are known. | absent or present |
| Foraging bias | With foraging bias, filling levels (if >0) are set to 1, making handling times independent of actual filling levels when bumblebees are exposed to the pesticide during OSR flowering period. | absent or present |

In the original foraging module, the model sorts the list of known foraging patches by their distances to the bee’s colony for each forager. In the presence of ‘ISI use’, foragers’ choice of where to collect nectar or pollen is influenced by the presence of conspecifics. Individuals’ known patches are first divided into two types: one type is where others are currently foraging, and the other type is the unoccupied patches. Then, both lists are sorted by according to the distance between each patch and the bee’s colony, and the lists are compiled so the occupied patches appear first in the compiled list (see the ‘Foraging_SortKnownPatchesListREP’ function in the extended code). The modified function is later called during both nectar and pollen foraging. Because of the way ISI use is coded, freshly emerged foragers are not affected by this parameter until they find at least two accessible patches (please note that foragers become experienced by acquiring knowledge of more and more foraging patches during the simulations). Thus, naïve individuals will deplete food sources within the grassland patch first, because these are always the closest to the nest, which assumption is consistent with empirical findings indicating that young workers fly much shorter distances than older ones (Gilgenreiner & Kurze 2024). Foragers can be attracted to the OSR patches by others only after they discovered those patches by themselves. Because of that, the contribution of ISI use to resource exploitation can be regarded as conservative in the extended model.

The foraging bias included in the extended model is based on a recent finding of Kárpáti et al. (2024). This study examined the peripheral olfactory detection of a floral blend and foraging behaviour of buff-tailed bumblebee workers exposed to a formulation containing acetamiprid, a neonicotinoid insecticide. The key result was that treated bumblebees were less likely to locate the floral blend source that was previously associated with syrup during chronic pesticide exposure, which led to a random choice between the control and reward-associated scent sources in the behavioural trials. Acetamiprid is frequently applied in OSR cultivations, particularly during the flowering period (Kathage et al. 2018; Dworzańska et al. 2020). As floral scent is generally associated with the presence of resources in flowers (Wright and Schiestl 2009; Knauer and Schiestl 2015), this finding suggests that exposed individuals could not correctly assess which presented scent source was rewarding. In BEE-STEWARD, floral scent cues are not represented explicitly; instead, the profitability of a food source is indicated by its resource filling level. To represent the above experimental finding, I implemented a ‘foraging bias’, in which affected bees choose among patches independently of filling level. This mechanism can be interpreted as an abstraction of reduced ability to use reward-associated cues during patch choice, rather than as a direct preference for any particular crop types. I modelled this effect by modifying the ‘Foraging_bestLayerREP’ function that reports the most profitable layer within the currently used flower patch based on minimal handling time. In the original model, handling time (‘handlingTimePerFlower_s’ function) for nectar foraging is determined by the time needed to travel to the next flower plus the time to test whether it contains nectar, divided by the filling level of the flower, plus the time needed to collect the nectar once a filled flower is found. During pollen foraging, handling time is calculated from the time needed to travel to the next flower multiplied by the number of flowers needed to be visited, divided by the filling level of the flower, plus the time needed to collect the pollen once a flower with pollen is found. In the presence of ‘foraging bias’, filling levels are set to 1 within foragers’ decision-making process, so handling times are not dependent on actual filling levels when bumblebees are exposed to a pesticide on crop patches. Still, only those patches are considered where the filling level > 0, i.e., flowering plants are present. I assumed that all bees are affected by the pesticide application on OSR patches irrespective of the patch that individuals are exploiting (as resources stored within the colonies can also be contaminated), but only during the OSR flowering period (i.e., from day 114 to day 136).

### Characteristics of the simulation runs

Simulations were run for 50 years in 50 iterations to determine if the default number of total hibernating queens (set to 500) is appropriate for the model and from which year the population can be assumed to be in equilibrium (Hui 2006; Fig. S1A). Based on the results, I took year 10 as the year when the bumblebee population was likely to reach equilibrium in the simulated landscape and thus subsequently analysed the data obtained in the 10^th^ year. The initial number of hibernating queens was also 500 in all iterations. I ran the simulations 50 times in all combinations of ‘ISI use’ (absent or present) and ‘foraging bias’ (absent or present), totalling 200 simulations. Models were run as individual-based simulations under the ‘intermediate foraging mortality’ scenario. I also used the setting where males were not unlimited.

### Data preparation

From the simulation outputs, I obtained the following measures from the 10^th^ simulation year for subsequent analysis: the number of visits associated with nectar foraging on each patch between day 114 and day 136 (i.e., during the OSR flowering period), the number of visits related to pollen foraging on each patch between day 114 and day 136, the quantities of nectar and pollen collected and stored in all colonies during this period, the total number of adult workers and the number of colonies on day 149, and the total number of hibernating queens on day 365. The total number of adult workers and the number of colonies were used as proxies for the pollination capacity of the simulated bumblebee population, as this time point falls within the period of peak annual population size. The total number of hibernating queens was considered as a proxy for fitness. These approximations are consistent with those applied in a similar simulation in BEE-STEWARD by Knapp et al. (2019). From the patch visits data, I calculated the proportions of nectar and pollen visits for the close and far OSR patches, respectively. For the distribution of total nectar and pollen visits between day 114 and day 136, see Fig. S2.

### Statistical analysis

All calculations were performed using R 4.5.0 (R Core Team 2025) and RStudio 2025.05.0 (Posit Team 2025). Instead of applying frequentist hypothesis testing during the analysis, I evaluated the magnitude of differences between simulation runs with different parameter settings and reported parameter estimates with corresponding uncertainty following the recommendations of White et al. (2014). I fitted generalised linear models (GLMs) with a beta binomial error distribution using the Template Model Builder (‘glmmTMB’ R Package; Brooks et al. 2017) with ISI use (as a factor), foraging bias (as a factor), and their two-way interaction as explanatory variables on the proportion of nectar and pollen visits, separately on the close and far OSR patches. With this design, I could explore whether the effects of these two processes are independent, additive, multiplicative, or whether they mitigate each other. In the model fitted on the proportion of pollen visits on the close OSR patches, ISI use was also included in the dispersion formula to account for variance differences between its levels. As a decision framework for evaluating differences in these models, I calculated odds ratios (OR) with corresponding 95% confidence intervals (CIs) as measures of effect size. To assess the impact of ISI use and foraging bias on the amount of collected resources during the OSR flowering period, I fitted linear models (with Gaussian error distribution; LMs) on the amount of stored nectar and pollen. To investigate the potential consequences of these behavioural parameters on pollination capacity and fitness, I also fitted LMs on the total number of adult workers, number of colonies, and total number of hibernating queens. Model assumptions were checked by plot diagnosis using the ‘DHARMa’ R package (Hartig 2022). I calculated estimated marginal means (EMMs) and contrasts on the response scale with 95% CIs using the ‘emmeans’ R package (Lenth 2023). For LMs, effects were considered negligible when the 95% confidence interval of these contrasts included zero.

## Results

### Nectar visits on close and far OSR patches

In the baseline setting, in which neither ISI use nor foraging bias was included, bumblebees visited the close OSR patches in 26.8% [25.6, 28.1] of the nectar foraging bouts (Fig. 2a). Incorporating ISI use was associated with a higher estimated proportion of visits to these patches (37.6% [36.2, 39.0]; contrast – presence vs. absence: 10.8 [8.9, 12.7]), and the estimated odds of visiting close OSR patches for nectar were higher by 65% (OR = 1.65 [1.51, 1.80]). Including foraging bias alone also increased the estimated proportion, although the magnitude of the contrast was smaller (31.1% [29.7, 32.4]; contrast: 4.2 [2.4, 6.1]; OR = 1.23 [1.12, 1.35]). When both behavioural parameters were included, the estimated proportion was similar to the ‘only ISI use’ setting (37.6% [36.1, 39.1]; contrast to the ‘only ISI use’ setting: <-0.01 [-2.1, 1.9]; OR = 1 [0.92, 1.08]).

**Figure 2.**
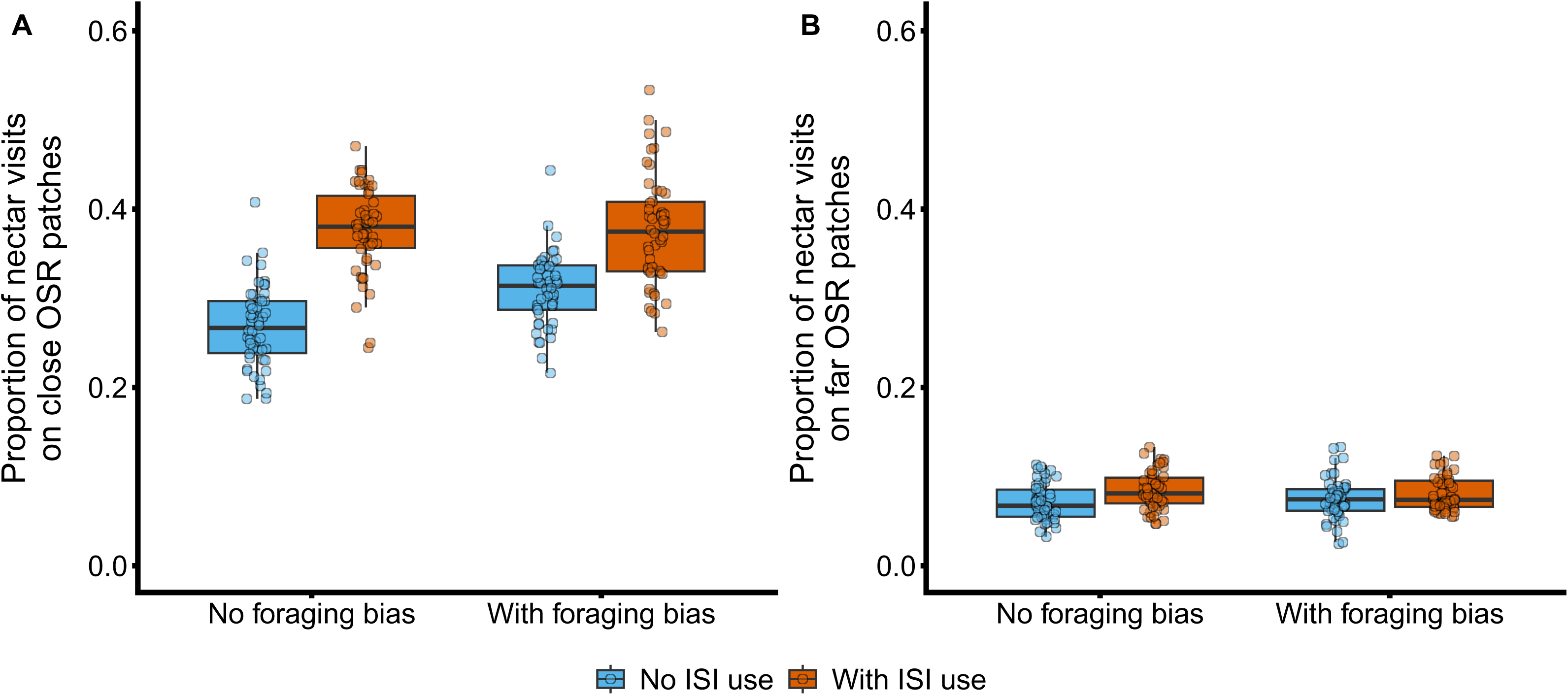
The proportion of nectar visits on the close (panel A) and far (panel B) OSR patches. Boxplots show the median and interquartile range; whiskers show values within 1.5-fold of the interquartile range, dots indicate individual values. The x-axis denotes the absence (‘No foraging bias’) or presence (‘With foraging bias’) of foraging bias. The filling colour of the boxplots indicate the absence (‘No ISI use’: teal) or presence of inadvertent social information (ISI) use (‘With ISI use’: orange). Alt text for Figure 2. Two-panel boxplot figure showing the proportion of nectar visits to oilseed rape (OSR) patches. Panel A represents close OSR patches and Panel B far OSR patches. The x-axis compares absence versus presence of foraging bias, and box colour indicates absence (teal) or presence (orange) of inadvertent social information (ISI) use. Boxplots display medians, interquartile ranges, whiskers (1.5× IQR), and individual data points.

On the far OSR patches, the estimated effects of ISI use and foraging bias followed a similar directional pattern but with smaller contrasts. In the absence of both behavioural parameters, the proportion of nectar visits was 7.1% [6.6, 7.7] (Fig. 2b). When ISI use was included, the estimated proportion increased to 8.5% [7.9, 9.1] (contrast – presence vs. absence: 1.3 [ 0.5, 2.2]; OR = 1.21 [1.08, 1.35]). When only foraging bias was present, the estimate was 7.4% [6.9, 8.0] (contrast: 0.3 [-0.5, 1.1]; OR = 1.05 [0.93, 1.17]). When both behavioural parameters were incorporated, the estimated proportion was similar to that observed with ISI use alone (8.1% [7.5, 8.7]; contrast to the ‘only ISI use’ setting: −0.3 [-1.2, 0.5]; OR = 0.96 [0.86, 1.07]).

### Pollen visits on close and far OSR patches

In the absence of ISI use and foraging bias, the proportion of pollen visits to the close OSR patches was 10.9% [9.3, 12.8] (Fig. 3a). When ISI use was included, the estimated proportion increased to 14.9% [13.3, 16.6] (contrast – presence vs. absence: 4.0 [1.5, 6.4]; OR = 1.43 [1.14, 1.78]). When only foraging bias was present, the proportion of pollen visits to the close OSR patches was almost three times higher than the baseline value (31.0% [28.4, 33.7]; contrast: 20.1 [16.9, 23.2]), corresponding to 3.66-fold higher odds of visitation (OR = 3.66 [2.95, 4.54]). When both behavioural parameters were included, the estimated proportion further increased to 34.3% [32.0, 36.6], corresponding to a contrast of 3.3 [<-0.1, 6.8] relative to the ‘only foraging bias’ setting (OR = 1.16 [0.99, 1.36]).

**Figure 3.**
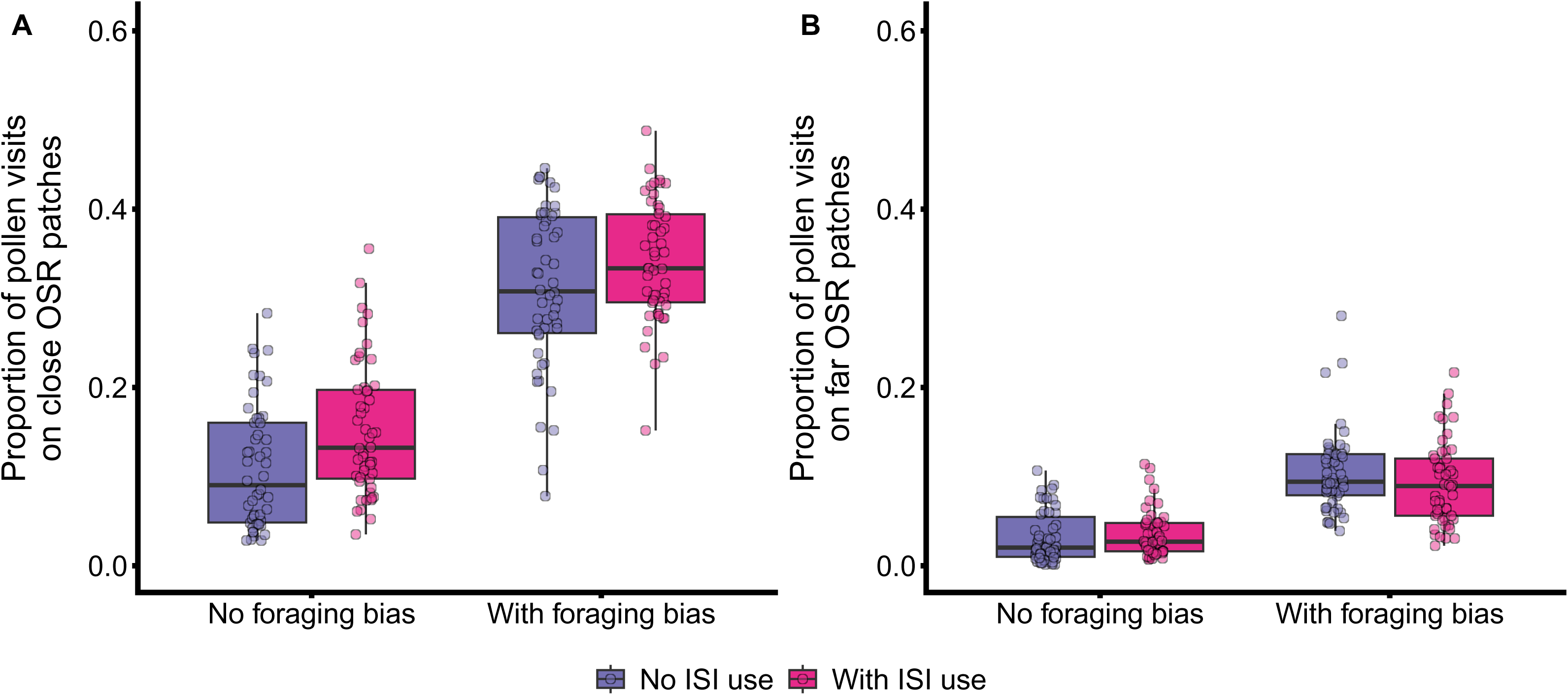
The proportion of pollen visits on the close (panel A) and far (panel B) OSR patches. Boxplots show the median and interquartile range (IQR). Whiskers denote the range of 1.5 times the IQR, while dots indicate individual data points. The x-axis denotes the absence (‘No foraging bias’) or presence (‘With foraging bias’) of foraging bias. The filling colour of the boxplots indicate the absence (‘No ISI use’: purple) or presence of ISI use (‘With ISI use’: magenta). Alt text for Figure 3. Two-panel boxplot figure showing the proportion of pollen visits to oilseed rape (OSR) patches. Panel A represents close OSR patches and Panel B far OSR patches. The x-axis compares absence versus presence of foraging bias, and box colour indicates absence (purple) or presence (magenta) of ISI use. Boxplots display medians, interquartile ranges, whiskers (1.5× IQR), and individual data points.

On the far OSR patches, the estimated proportion of pollen visits was 2.9% [2.4, 3.6] in the absence of both behavioural parameters. When only ISI use was present, the estimated proportion was 3.8% [3.1, 4.6] (contrast: 0.8 [-0.1, 1.7]; OR = 1.3 [0.98, 1.71]). When foraging bias was included, the proportion increased to 10.6 % [9.4, 11.9] (contrast – presence vs. absence: 7.6 [6.3, 9.0]; Fig. 3b), and the odds of visiting far OSR patches for pollen were 3.9 times higher (OR = 3.9 [3.05, 5.0]). When both behavioural parameters were included, the estimate was 9.1% [8.0, 10.3], corresponding to a contrast of −1.4 [−3.1, <0.01] relative to the ‘only foraging bias’ setting (OR = 0.85 [0.7, 1.03]).

### Resources collected during the OSR flowering period

The amount of nectar stored in the baseline setting was 146 mL [127, 165]. When ISI use was included, the estimated amount was 170 mL [152, 189] (contrast – presence-absence: 24.8 [-1.8, 51.3]), and when only foraging bias was present, the estimate was 169 mL [150, 188] (contrast: 23.6 [-3, 50.1]). When both behavioural parameters were included, the estimated amount was 167 mL [148, 186] (contrast relative to the baseline setting: 21.5 [-5.1, 48.1]).

The amount of pollen stored in the baseline setting was 60.8 g [54.6, 67.1]. When ISI use was present, the estimated amount was 70.9 g [64.6, 77.1] (contrast – presence vs. absence: 10 [1.2, 18.9]). When only foraging bias was included, the estimate was 66.5 g [60.2, 72.8] (contrast: 5.7 [-3.2, 14.6]). When both parameters were included, the estimated amount of pollen stored was 63.7 g [57.5, 70.0] (contrast relative to the baseline setting: 2.9 [-5.97, 11.8]).

### Consequences on pollination capacity and fitness

In the baseline setting, the simulated landscape contained a total of 373 adult workers [336, 409], 37.3 colonies [33.7, 40.9], and 452 hibernating queens [420, 485]. The estimated numbers of adult workers and hibernating queens were similar across scenarios that included ISI use, foraging bias, or both behavioural parameters. When both ISI use and foraging bias were present, the estimated number of colonies was slightly higher than in the baseline setting (contrast relative to the baseline setting: 6.1 [1, 11.2]; Table 2).

**Table 2.** Estimated marginal means (EMMs) of the proxy measures for pollination capacity and fitness in different combinations of the investigated behavioural parameters. EMMs and contrasts are on the response scale and reported with their corresponding 95% confidence intervals (in square brackets). All values were estimated from the fitted LMs.

| Simulation measures | Baseline setting | With ISI use (presence vs. absence contrast) | With foraging bias (presence vs. absence contrast) | With both ISI use and foraging bias (contrast relative to the baseline setting) |
| --- | --- | --- | --- | --- |
| Total number of adult workers | 373 [336, 409] | 420 [384, 457]<br>(47.8 [-4, 99.5]) | 390 [353, 426]<br>(17 [-34.7, 68.8]) | 384 [348, 421]<br>(11.8 [-40, 63.5]) |
| Number of colonies | 37.3 [33.7, 40.9] | 41.3 [37.7, 44.9]<br>(4 [-1.1, 9.12]) | 40.5 [36.9, 44.1]<br>(3.2 [-1.9, 8.3]) | 43.4 [39.8, 47.0]<br>(6.1 [1, 11.2]) |
| Total number of hibernating queens | 452 [420, 485] | 443 [410, 475]<br>(-9.6 [-55.6, 36.4]) | 439 [407, 472]<br>(-12.7 [-58.7, 33.3]) | 458 [426, 491]<br>(6 [-40, 52]) |

## Discussion

Understanding how ecological traits shape the interaction of non-*Apis* bees with heterogeneous landscapes remains an important challenge (Raine and Rundlöf 2024). By incorporating inadvertent social information (ISI) use and a recently documented foraging bias into the BEE-STEWARD model, I found that patch visitation patterns of buff-tailed bumblebees shifted relative to the original model. Specifically, ISI use enhanced nectar visitation to both close and far OSR patches and also increased pollen visitation to the close crop patches. Foraging bias primarily increased pollen visitation to all OSR patches and also increased nectar visitation to close OSR patches. I have found no evidence of an interactive effect between the two parameters on crop patch visitations. The observed shifts in patch visitation did not alter the amount of resource collected or induced major changes in colony growth parameters and reproductive performance. Overall, these simulations show that in a crop patch-dominated landscape where the bumblebee population is at equilibrium, both behavioural parameters can increase foragers’ resource collection rate from crop patches, despite the availability of alternative food sources during crop bloom (also in Carturan et al. 2023). The presented model does not explicitly simulate pesticide residues, intake, or dose accumulation, and therefore does not provide quantitative estimates of pesticide exposure levels. Instead, it quantifies how ISI use and foraging biases shape visitation patterns across different patch types. Consequently, the model identifies two previously overlooked behavioural mechanisms that can substantially alter spatial foraging patterns of bumblebees in agricultural landscapes.

Bumblebees are central place foragers (Heinrich 1979), and most recent models have simulated their foraging behaviour as primarily depending on the distance to nest and floral resource quality (Olsson et al. 2015; Häussler et al. 2017; Blasi et al. 2022; Carturan et al. 2023; Lonsdorf et al. 2024). These variables are regarded as key drivers of bumblebees’ foraging decisions because they can accurately predict resource partitioning especially at larger spatial scales and are well-supported by empirical data (Lihoreau et al. 2011; Pope and Jha 2018; Fragoso et al. 2021). Nevertheless, findings of this study indicate that considering foraging decisions at a finer resolution and taking additional behavioural pathways into account can be crucial for accurately estimating patch visitation patterns in bumblebees.

Conspecific attraction among foragers has been proposed to play a key role in the exploitation of patchily distributed resources (Dubois et al. 2021), whereas experimentally documented foraging biases have not yet been integrated into similar modelling frameworks. Although these behavioural parameters had limited effects on total resource intake and colony demographics, both altered the origin of the collected resources, indicating subtle changes in the realised foraging domain. Such shifts in resource use may have important ecological consequences by changing the relative contribution of different habitat types to colony resource acquisition. Previous empirical work has shown that foraging patterns can influence pollen-based pesticide risk in *B. terrestris* (Knapp et al. 2023), but assessing such effects was beyond the scope of the present study.

Social interactions can influence foraging decisions in bumblebees, yet the extent to which ISI use influences the visitation rates across different food patches remains poorly understood (unlike in honeybees; Hasenjager et al. 2024). In the context of nectar foraging, ISI use emerged as an influential factor shaping spatial foraging patterns by attracting conspecifics to crop patches. However, its effect was spatially limited but additive with foraging bias on pollen visitation rates. I modelled ISI use as a density-dependent response (effectively a ‘copy-the-majority’ rule) rather than a state-dependent ‘copy-when-uncertain’ strategy (as suggested by Smolla et al. 2016), because formalising ‘uncertainty’ in a spatially explicit context remains methodologically challenging. Furthermore, simple attraction to conspecific density is often a proximate mechanism driving social aggregation, even if the ultimate function aligns with reducing uncertainty (Leadbeater and Chittka 2007).

Nevertheless, this implementation corresponds to empirical findings indicating that social and personal information interact synergistically, wherein individuals do not prioritize social cues above direct experience (Leadbeater and Florent 2014; Smolla et al. 2016). I predicted that bumblebees would alter their patch visitation patterns and exploit the available food sources less efficiently in the presence of foraging bias, and this effect may be further amplified by the presence of ISI use via attracting conspecifics to suboptimal resource patches. However, the interplay between these two behavioural parameters was negligible, likely because the simulated landscape was coarse-grained and included a proximal grassland patch that could fully support all colonies throughout the year. Due to this landscape structure, foragers may not need to copy others to locate additional resource patches, regardless of whether a foraging bias is present. Consequently, the influence of ISI use could be substantially larger in more heterogeneous, fine-grained environments, and may interact more strongly with foraging biases, particularly if resources in the nesting habitat become insufficient during crop bloom.

Simulations indicated that the incorporated foraging bias was a major driver of pollen visitation rates to crop patches, regardless of patch distance and outweighing the influence of ISI use. I propose that this bias did not directly cause bees to shift from the grassland patch to OSR patches. Instead, it reduced their preference for more rewarding resources and thereby weakened discrimination among available food sources within the grassland patch, where multiple floral resources coexist. At the landscape scale, this also diverted some foragers from the grassland patch to OSR patches. As a result, the distribution of exploited patches more closely reflected the distribution of available patches within the colony’s foraging range. This also suggests that in landscapes containing multiple semi-natural habitat patches, the effect of such a foraging bias may be less pronounced compared to that observed in these simulations. Field studies have demonstrated that some pesticide-induced foraging biases can reduce colony-level resource collection in bumblebees (e.g., Gill and Raine 2014), but their direct effects on spatial visitation patterns and landscape-level resource exploitation have rarely been examined.

Foraging biases in pollinators may emerge through multiple mechanisms due to exposure to agrochemicals. First, insecticides can directly affect chemosensory orientation (Kárpáti et al. 2024), resulting in suboptimal choices between rewarding and unrewarding patches. Alternatively, such substances can impair spatial and chemical learning (Samuelson et al. 2016; Muth et al. 2019), cognitive abilities (Andrione et al. 2016; Gray et al. 2024), floral preference (Gill and Raine 2014; Phelps et al. 2018), or reduce foraging motivation (Siviter et al. 2021; Ohlinger et al. 2022; Siviter and Muth 2022), all of which may indirectly lead to foraging decisions that are similar to the modelled foraging bias.

Patch choice decisions related to nectar and pollen visits differ in the Bumble-BEEHAVE model incorporated in BEE-STEWARD. The calculation of handling time for nectar foragers follows the model of Harder (1983) and energetic efficiency of exploiting a patch (i.e., energy gained compared to the energy needed to exploit a given layer) plays a crucial role in determining the most profitable food source. However, pollen foraging is modelled to maximise foraging rate. In addition, the priority of nectar and pollen foraging is also different in the model, and individuals always search for nectar if the stimulus for this task in the bee’s colony is higher than the bee’s associated threshold, regardless of the stimulus-threshold relationship of the other tasks (including pollen foraging). This difference may, at least partially, explain why the examined behavioural parameters had distinct impacts on the nectar and pollen visits in the simulations.

In conclusion, this study shows that previously overlooked aspects of foraging behaviour can alter the visitation rates to available resource patches in buff-tailed bumblebees, which is consistent with field observations showing that mass-flowering crops can competitively dilute pollinator densities in agricultural landscapes (Holzschuh et al. 2011; Riggi et al. 2024). In particular, simulation results imply that both ISI use and an experimentally documented foraging bias may lead to more frequent visitation of crop patches, without changing overall resource collection. Such shifts in patch exploitation may have implications for pesticide exposure risk in real-world agricultural landscapes, although these processes were not explicitly modelled here. Future efforts should extend the BEE-STEWARD model to additional behavioural processes that may influence spatial foraging patterns and patch visitation. Incorporating pesticide concentrations in floral resources and their temporal dynamics, together with sublethal effects on colony performance (Stanley et al. 2015; Baron et al. 2017), would allow future studies to evaluate how behaviourally driven visitation patterns translate into exposure risk and demographic consequences.

## Acknowledgements

ZT was financially supported by the János Bolyai Research Scholarship of the Hungarian Academy of Sciences (MTA, BO/00634/21/8) and by the Scientific Patronage Grant of the Ministry for Innovation and Technology from the source of the National Research, Development and Innovation Fund (NKFIH, MEC_N_149199).

## Author contribution statement

The author confirms sole responsibility for the following: conceptualisation, methodology, data curation, formal analysis, investigation, validation, visualisation, and writing.

## Data availability statement

Input files for the extended BEE-STEWARD model, R script for the analysis of model outputs, and data supporting the results are archived and available at Figshare (https://figshare.com/s/acee9613adfc7757267b).

## Ethics approval

Not applicable.

## Declaration of generative AI and AI-assisted technologies in the writing process

During the preparation of this work, the author used the GPT-4o and GPT-5 models (OpenAI Inc., USA) to assist writing and improve the readability and language of the manuscript. After using this product, the author reviewed and edited the content as needed and take full responsibility for the content of the published article.

## References

Andrione M, Vallortigara G, Antolini R, Haase A, 2016. Neonicotinoid-induced impairment of odour coding in the honeybee. Sci Rep 6: 38110.

Arce AN, Ramos Rodrigues A, Yu J, Colgan TJ, Wurm Y et al., 2018. Foraging bumblebees acquire a preference for neonicotinoid-treated food with prolonged exposure. Proc R Soc B 285: 20180655.

Avarguès-Weber A, Chittka L, 2014. Observational conditioning in flower choice copying by bumblebees (*Bombus terrestris*): influence of observer distance and demonstrator movement. PLoS ONE 9: e88415.

Avarguès-Weber A, Lachlan R, Chittka L, 2018. Bumblebee social learning can lead to suboptimal foraging choices. Anim Behav 135: 209–214.

Banks SC, Lindenmayer DB, Wood JT, McBurney L, Blair D et al., 2013. Can individual and social patterns of resource use buffer animal populations against resource decline? PLoS ONE 8: e53672.

Baron GL, Jansen VA, Brown MJ, Raine NE, 2017. Pesticide reduces bumblebee colony initiation and increases probability of population extinction. Nat Ecol Evol 1: 1308–1316.

Baude M, Dajoz I, Danchin E, 2008. Inadvertent social information in foraging bumblebees: effects of flower distribution and implications for pollination. Anim Behav 76: 1863–1873.

Becher MA, Grimm V, Knapp J, Horn J, Twiston-Davies G et al., 2016. BEESCOUT: A model of bee scouting behaviour and a software tool for characterizing nectar/pollen landscapes for BEEHAVE. Ecol Modell 340: 126–133.

Becher MA, Twiston-Davies G, Penny TD, Goulson D, Rotheray EL et al., 2018. Bumble-BEEHAVE: A systems model for exploring multifactorial causes of bumblebee decline at individual, colony, population and community level. J Appl Ecol 55: 2790–2801.

Blasi M, Clough Y, Jönsson AM, Sahlin U, 2022. A model of wild bee populations accounting for spatial heterogeneity and climate-induced temporal variability of food resources at the landscape level. Ecol Evol 12: e9014.

Brooks ME, Kristensen K, Van Benthem KJ, Magnusson A, Berg CW et al., 2017. glmmTMB balances speed and flexibility among packages for zero-inflated generalized linear mixed modeling. R J 9: 378–400.

Byrne FJ, Visscher PK, Leimkuehler B, Fischer D, Grafton-Cardwell EE, et al., 2014. Determination of exposure levels of honey bees foraging on flowers of mature citrus trees previously treated with imidacloprid. Pest Manag Sci 70: 470–482.

Carturan BS, Siewe N, Cobbold CA, Tyson RC, 2023. Bumble bee pollination and the wildflower/crop trade-off: When do wildflower enhancements improve crop yield? Ecol Modell 484: 110447.

Coolen I, Dangles O, Casas J, 2005. Social learning in noncolonial insects? Curr Biol 15: 1931–1935.

Crall JD, De Bivort BL, Dey B, Ford Versypt AN, 2019. Social buffering of pesticides in bumblebees: agent-based modeling of the effects of colony size and neonicotinoid exposure on behavior within nests. Front Ecol Evol 7: 51.

Cruz A, Heinemans M, Marquez C, Moita MA, 2020. Freezing displayed by others is a learned cue of danger resulting from co-experiencing own freezing and shock. Curr Biol 30: 1128–1135.

Curk T, Rast W, Portas R, Kohles J, Shatumbu G et al., 2025. Advantages and disadvantages of using social information for carcass detection – A case study using white-backed vultures. Ecol Modell 499: 110941.

Danchin É, Giraldeau L-A, Valone TJ, Wagner RH, 2004. Public information: from nosy neighbors to cultural evolution. Science 305: 487–491.

Dawson EH, Chittka L, 2014. Bumblebees (*Bombus terrestris*) use social information as an indicator of safety in dangerous environments. Proc R Soc B 281: 20133174.

Donaldson R, Finn H, Bejder L, Lusseau D, Calver M, 2012. The social side of human–wildlife interaction: wildlife can learn harmful behaviours from each other. Anim Conserv 15: 427–435.

Dubois T, Pasquaretta C, Barron AB, Gautrais J, Lihoreau M, 2021. A model of resource partitioning between foraging bees based on learning. PLoS Comput Biol 17: e1009260.

Dworzańska D, Moores G, Zamojska J, Strażyński P, Węgorek P, 2020. The influence of acetamiprid and deltamethrin on the mortality and behaviour of honeybees (*Apis mellifera carnica* Pollman) in oilseed rape cultivations. Apidologie 51: 1143–1154.

Fragoso FP, Jiang Q, Clayton MK, Brunet J, 2021. Patch selection by bumble bees navigating discontinuous landscapes. Sci Rep 11: 8986.

Gil MA, Hein AM, 2017. Social interactions among grazing reef fish drive material flux in a coral reef ecosystem. P Natl Acad Sci USA 114: 4703–4708.

Gil MA, Baskett ML, Schreiber SJ, 2019. Social information drives ecological outcomes among competing species. Ecology 100: e02835.

Gil MA, Hein AM, Spiegel O, Baskett ML, Sih A, 2018. Social information links individual behavior to population and community dynamics. Trends Ecol Evol 33: 535–548.

Gill RJ, Raine NE, 2014. Chronic impairment of bumblebee natural foraging behaviour induced by sublethal pesticide exposure. Funct Ecol 28: 1459–1471.

Gray LK, Hulsey M, Siviter H, 2024. A novel insecticide impairs bumblebee memory and sucrose responsiveness across high and low nutrition. Roy Soc Open Sci 11: 231798.

Gruber T, Luncz L, Mörchen J, Schuppli C, Kendal RL et al., 2019. Cultural change in animals: a flexible behavioural adaptation to human disturbance. Palgrave Commun 5: 1–9.

Grüter C, Leadbeater E, 2014. Insights from insects about adaptive social information use. Trends Ecol Evol 29: 177–184.

Harder LD, 1983. Flower handling efficiency of bumble bees: morphological aspects of probing time. Oecologia 57: 274–280.

Hartig F, 2022. DHARMa: Residual Diagnostics for Hierarchical (Multi-Level/Mixed) Regression Models. R package version 0.4.6.

Hasenjager MJ, Hoppitt W, Cunningham-Eurich I, Franks VR, Leadbeater E, 2024. Coupled information networks drive honeybee (*Apis mellifera*) collective foraging. J Anim Ecol 93: 71–82.

Häussler J, Sahlin U, Baey C, Smith HG, Clough Y, 2017. Pollinator population size and pollination ecosystem service responses to enhancing floral and nesting resources. Ecol Evol 7: 1898–1908.

Heinrich B, 2004. Bumblebee economics. Cambridge, MA: Harvard University Press.

Holzschuh A, Dormann CF, Tscharntke T, Steffan-Dewenter I, 2011. Expansion of mass-flowering crops leads to transient pollinator dilution and reduced wild plant pollination. Proc R Soc B 278: 3444–3451.

Hui C, 2006. Carrying capacity, population equilibrium, and environment’s maximal load. Ecol Modell 192: 317–320.

Hutchinson LA, Oliver TH, Breeze TD, Bailes EJ, Brünjes L et al., 2021. Using ecological and field survey data to establish a national list of the wild bee pollinators of crops. Agric Ecosyst Environ 315: 107447.

Jones PL, Ryan MJ, Chittka L, 2015. The influence of past experience with flower reward quality on social learning in bumblebees. Anim Behav 101: 11–18.

Kárpáti Z, Szelényi MO, Tóth Z, 2024. Exposure to an insecticide formulation alters chemosensory orientation, but not floral scent detection, in buff-tailed bumblebees (*Bombus terrestris*). Sci Rep 14: 14622.

Kathage J, Castañera P, Alonso-Prados JL, Gómez-Barbero M, Rodríguez-Cerezo E, 2018. The impact of restrictions on neonicotinoid and fipronil insecticides on pest management in maize, oilseed rape and sunflower in eight European Union regions. Pest Manag Sci 74: 88–99.

Kawaguchi LG, Ohashi K, Toquenaga Y, 2006. Do bumble bees save time when choosing novel flowers by following conspecifics? Funct Ecol 20: 239–244.

Kessler SC, Tiedeken EJ, Simcock KL, Derveau S, Mitchell J et al., 2015. Bees prefer foods containing neonicotinoid pesticides. Nature 521: 74–76.

Knapp JL, Becher MA, Rankin CC, Twiston-Davies G, Osborne JL, 2019. *Bombus terrestris* in a mass-flowering pollinator-dependent crop: A mutualistic relationship? Ecol Evol 9: 609–618.

Knapp JL, Nicholson CC, Jonsson O, de Miranda JR, Rundlöf M, 2023. Ecological traits interact with landscape context to determine bees’ pesticide risk. Nat Ecol Evol 7: 547–556.

Knauer AC, Schiestl FP, 2015. Bees use honest floral signals as indicators of reward when visiting flowers. Ecol Lett 18: 135–143.

Kokko H, Sutherland WJ, 2001. Ecological traps in changing environments: ecological and evolutionary consequences of a behaviourally mediated Allee effect. Evol Ecol Res 3: 603–610.

Laidre ME, 2010. How rugged individualists enable one another to find food and shelter: field experiments with tropical hermit crabs. Proc R Soc B 277: 1361–1369.

Lämsä J, Kuusela E, Tuomi J, Juntunen S, Watts PC, 2018. Low dose of neonicotinoid insecticide reduces foraging motivation of bumblebees. Proc R Soc B 285: 20180506.

Laundré JW, Hernández L, Ripple WJ, 2010. The landscape of fear: ecological implications of being afraid. Open Ecol J 3: 1–7.

Leadbeater E, 2015. What evolves in the evolution of social learning? J Zool 295:4–11.

Leadbeater E, Chittka L, 2009. Bumble-bees learn the value of social cues through experience. Biol Lett 5: 310–312.

Leadbeater E, Florent C, 2014. Foraging bumblebees do not rate social information above personal experience. Behav Ecol Sociobiol 68: 1145–1150.

Lenth R, 2023. emmeans: Estimated marginal means, aka least-squares means. R package version 1.8.6.

Lihoreau M, Chittka L, Raine NE, 2011. Trade-off between travel distance and prioritization of high-reward sites in traplining bumblebees. Funct Ecol 25: 1284–1292.

Lonsdorf EV, Rundlöf M, Nicholson CC, Williams NM, 2024. A spatially explicit model of landscape pesticide exposure to bees: Development, exploration, and evaluation. Sci Total Environ 908: 168146.

Loukola OJ, Solvi C, Coscos L, Chittka L, 2017. Bumblebees show cognitive flexibility by improving on an observed complex behavior. Science 355: 833–836.

Martín J, Luque-Larena JJ, López P, 2006. Collective detection in escape responses of temporary groups of Iberian green frogs. Behav Ecol 17: 222–226.

Muth F, Leonard AS, 2019. A neonicotinoid pesticide impairs foraging, but not learning, in free-fying bumblebees. Sci Rep 9: 4764.

Muth F, Francis JS, Leonard AS, 2019. Modality-specific impairment of learning by a neonicotinoid pesticide. Biol Lett 15: 20190359.

Ohlinger BD, Schürch R, Durzi S, Kietzman PM, Silliman MR et al., 2022. Honey bees (Hymenoptera: Apidae) decrease foraging but not recruitment after neonicotinoid exposure. J Insect Sci 22: 16.

Olsson O, Bolin A, Smith HG, Lonsdorf EV, 2015. Modeling pollinating bee visitation rates in heterogeneous landscapes from foraging theory. Ecol Modell 316: 133–143.

Parejo D, Avilés JM, 2016. Social information use by competitors: resolving the enigma of species coexistence in animals? Ecosphere 7: e01295.

Pays O, Beauchamp G, Carter AJ, Goldizen AW, 2013. Foraging in groups allows collective predator detection in a mammal species without alarm calls. Behav Ecol 24: 1229–1236.

Perrot T, Bretagnolle V, Acar N, Febvret V, Matejicek A et al., 2024. Bees improve oil quality of oilseed rape. Basic Appl Ecol 76: 41–49.

Phelps JD, Strang CG, Gbylik-Sikorska M, Sniegocki T, Posyniak A et al., 2018. Imidacloprid slows the development of preference for rewarding food sources in bumblebees (Bombus impatiens). Ecotoxicology 27: 175–187.

Pohorecka K, Skubida P, Miszczak A, Semkiw P, Sikorski P et al., 2012. Residues of neonicotinoid insecticides in bee collected plant materials from oilseed rape crops and their effect on bee colonies. J Apic Sci 56: 115.

Pope NS, Jha S, 2018. Seasonal food scarcity prompts long-distance foraging by a wild social bee. Am Nat 191: 45–57.

Posit Team RStudio, 2024. Integrated Development Environment for R. Posit Software, PBC, Boston, MA.

Pouca CV, Heinrich D, Huveneers C, Brown C, 2020. Social learning in solitary juvenile sharks. Anim Behav 159: 21–27.

R Core Team R, 2023. A language and environment for statistical computing. R Foundation for Statistical Computing, Vienna, Austria. https://www.R-project.org/

Raine NE, Rundlöf M, 2024. Pesticide exposure and effects on non-*Apis* bees. Annu Rev Entomol 69: 551–576.

Riggi LG, Raderschall CA, Fijen TPM, Scheper J, Smith HG et al., 2024. Early-season mass-flowering crop cover dilutes wild bee abundance and species richness in temperate regions: A quantitative synthesis. J Appl Ecol 61: 452–464.

Samuelson EE, Chen-Wishart ZP, Gill RJ, Leadbeater E, 2016. Effect of acute pesticide exposure on bee spatial working memory using an analogue of the radial-arm maze. Sci Rep 6: 38957.

Schmidt KA, Johansson J, Betts MG, 2015. Information-mediated Allee effects in breeding habitat selection. Am Nat 186(6): e162–e171.

Seppänen JT, Forsman JT, Mönkkönen M, Thomson RL, 2007. Social information use is a process across time, space, and ecology, reaching heterospecifics. Ecology 88: 1622–1633.

Sigaud M, Merkle JA, Cherry SG, Fryxell JM, Berdahl A et al., 2017. Collective decision-making promotes fitness loss in a fusion-fission society. Ecol Lett 20:33–40.

Siviter H, Johnson AK, Muth F, 2021. Bumblebees exposed to a neonicotinoid pesticide make suboptimal foraging decisions. Environ Entomol 50: 1299–1303.

Siviter H, Muth F, 2022. Exposure to the novel insecticide flupyradifurone impairs bumblebee feeding motivation, learning, and memory retention. Environ Pollut 307: 119575.

Smolla M, Alem S, Chittka L, Shultz S, 2016. Copy-when-uncertain: bumblebees rely on social information when rewards are highly variable. Biol Lett 12: 20160188.

Stanley DA, Garratt MP, Wickens JB, Wickens VJ, Potts SG et al., 2015. Neonicotinoid pesticide exposure impairs crop pollination services provided by bumblebees. Nature 528: 548–550.

Tóth Z, Jaloveczki B, Tarján G, 2020. Diffusion of Social Information in Non-grouping Animals. Front Ecol Evol 8: 586058.

Twiston-Davies G, Becher MA, Osborne JL, 2021. BEE-STEWARD: A research and decision-support software for effective land management to promote bumblebee populations. Methods Ecol Evol 12: 1809–1815. Ward LT, Hladik ML, Guzman A, Winsemius S, Bautista A, Kremen C, Mills NJ, 2022. Pesticide exposure of wild bees and honey bees foraging from field border flowers in intensively managed agriculture areas. Sci Total Environ 831: 154697.

Webster MM, Laland KN, 2017. Social information use and social learning in non-grouping fishes. Behav Ecol 28: 1547–1552.

White JW, Rassweiler A, Samhouri JF, Stier AC, White C, 2014. Ecologists should not use statistical significance tests to interpret simulation model results. Oikos 123: 385–388.

Wilensky U, 1999. NetLogo. Center for Connected Learning and Computer-Based Modeling. Northwestern University, Evanston, IL. http://ccl.northwestern.edu/netlogo

Wilson MW, Ridlon AD, Gaynor KM, Gaines SD, Stier AC et al., 2020. Ecological impacts of human-induced animal behaviour change. Ecol Lett 23: 1522–1536.

Worden BD, Papaj DR, 2005. Flower choice copying in bumblebees. Biol Lett 1: 504–507.

Wray MK, Klein BA, Seeley TD, 2012. Honey bees use social information in waggle dances more fully when foraging errors are more costly. Behav Ecol 23: 125–131.

Wright GA, Schiestl FP, 2009. The evolution of floral scent: the influence of olfactory learning by insect pollinators on the honest signalling of floral rewards. Funct Ecol 23: 841–851.

Yokoi T, Goulson D, Fujisaki K, 2007. The use of heterospecific scent marks by the sweat bee *Halictus aerarius*. Sci Nat 94: 1021–1024.

