## Supplementary material for "Modelling the interplay between inadvertent social information use and a pesticide-induced foraging bias in bumblebees’ crop visitation": Figure S1

Zoltán Tóth^1^

^1^Department of Zoology, Plant Protection Institute, HUN-REN Centre for Agricultural Research, Budapest, Hungary

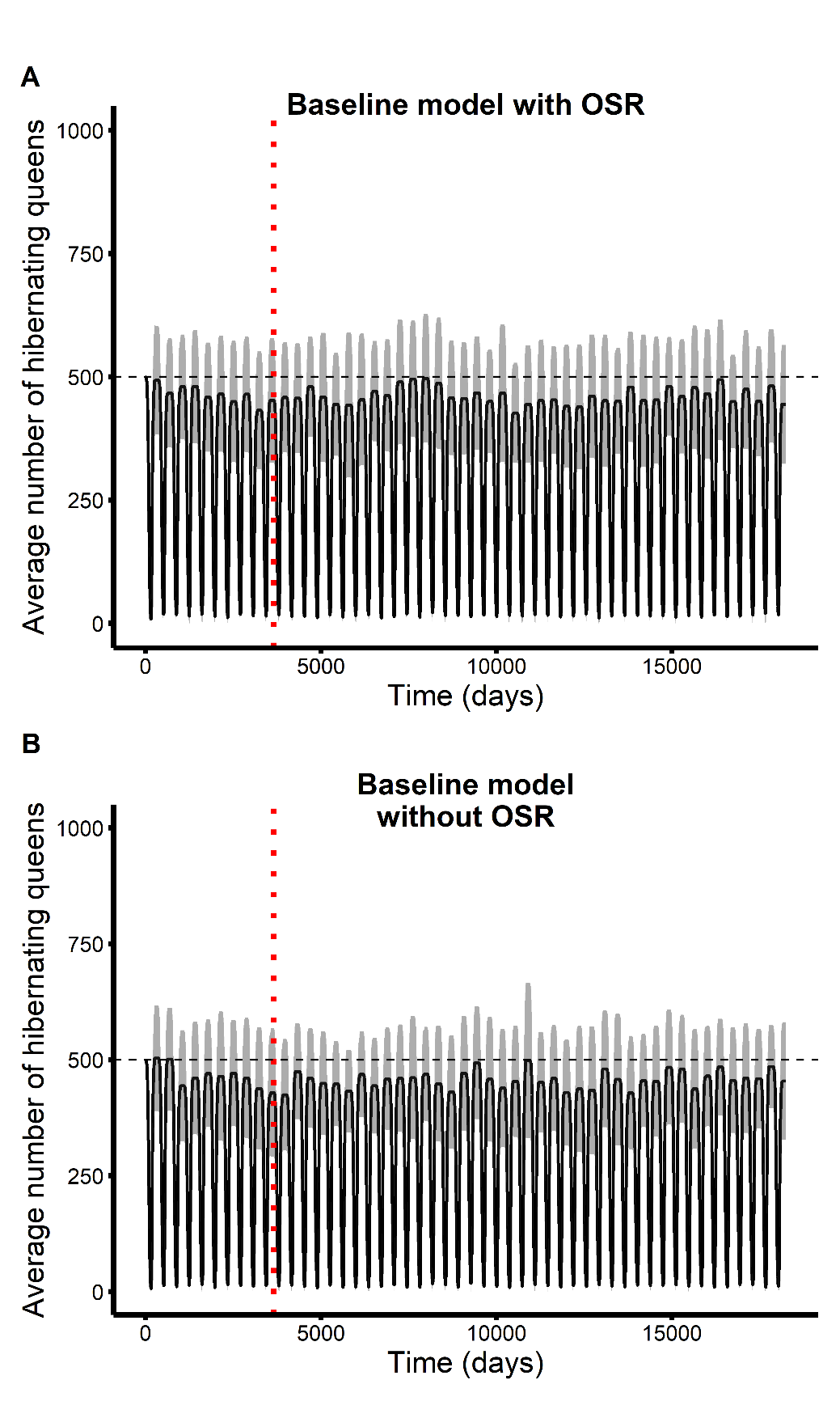

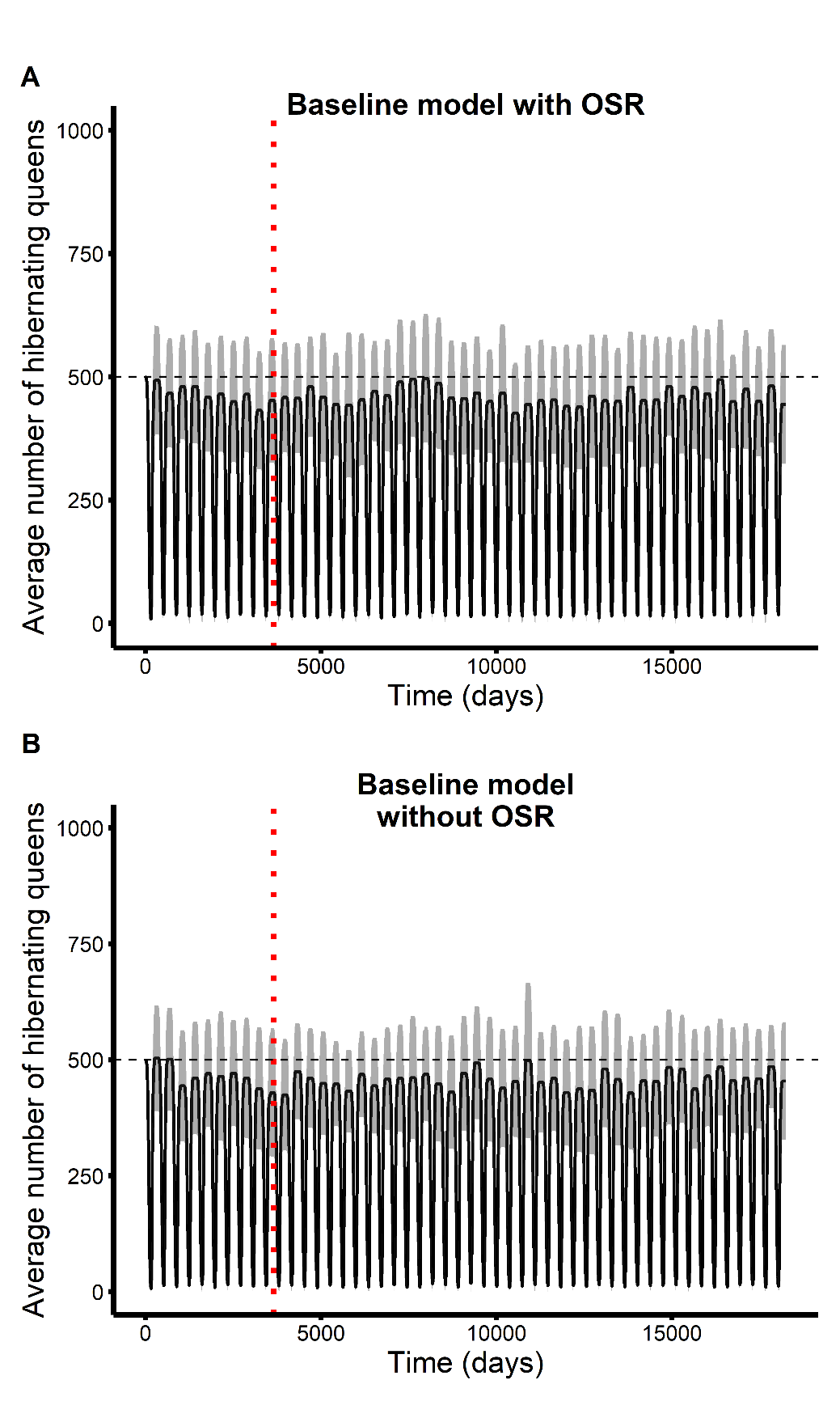

Figure S1. The average number of hibernating queens ± SD (shaded) across 50 iterations over 50 years in the presence (A) and absence of OSR patches (B) in the simulated landscape. The vertical dotted red line indicates the end of the 10^th^ year, from which the bumblebee population was considered likely to be in equilibrium. The horizontal dashed black line denotes the default setting of 500 initial hibernating queens at the start of each iteration. Simulations were run in the ‘baseline’ scenario, i.e., using the original model.

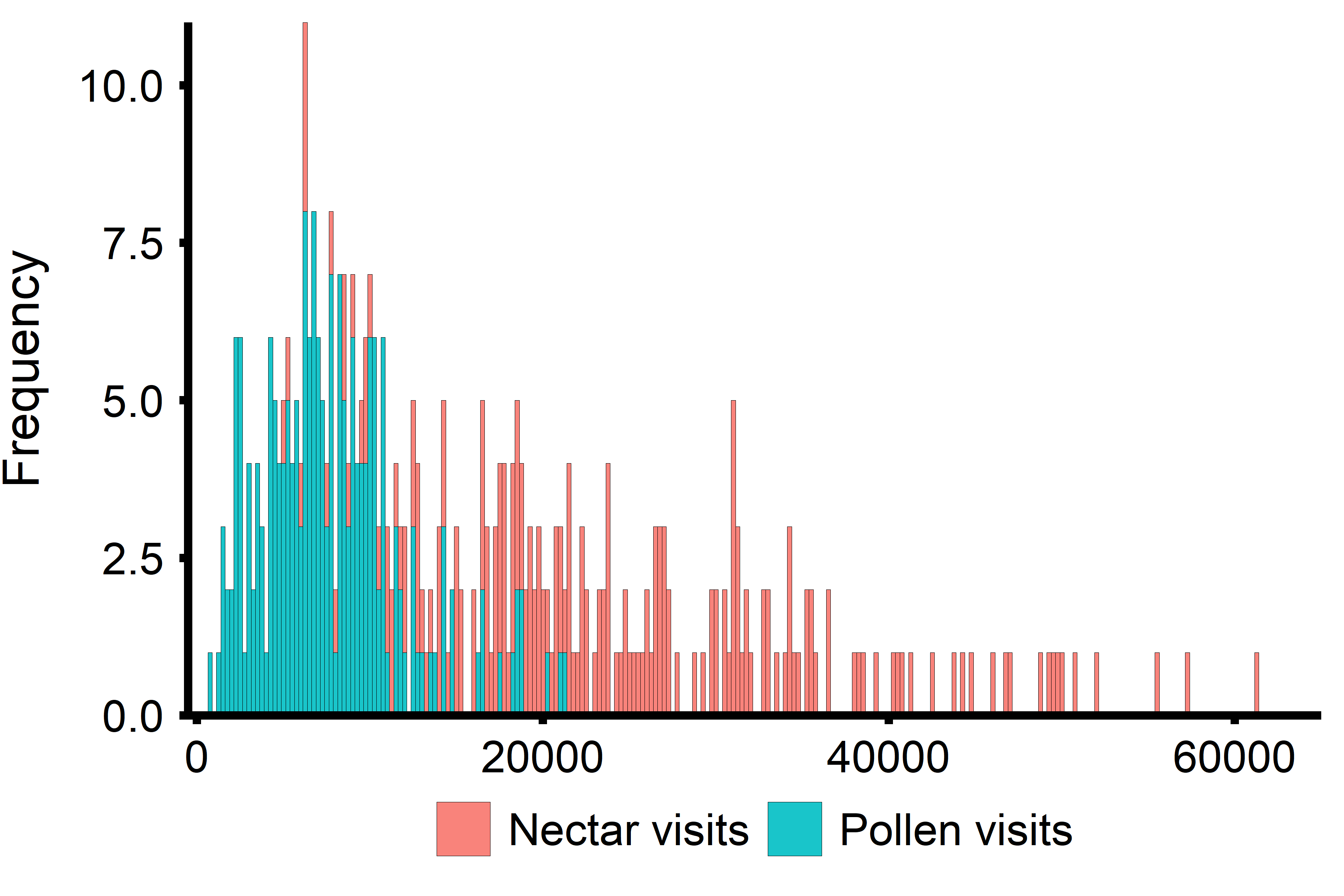

Figure S2. Distributions of the total number of nectar (in reddish-pink) and pollen (in cyan blue) visits between days 114–136. The number of nectar visits ranged from 5092 to 61179, whereas the number of pollen visits ranged from 656 to 21219. Simulations were run under baseline settings in a landscape containing OSR patches (*N*=30 iterations), and the reported values for nectar and pollen visits are from the 10^th^ simulation year.
